# Inhibitory Effect of *Bacillus velezensis* on Biofilm Formation by *Streptococcus mutans*

**DOI:** 10.1101/313965

**Authors:** Yesol Yoo, Dong-Ho Seo, Hyunjin Lee, Young-Do Nam, Myung-Ji Seo

**Affiliations:** Department of Bioengineering and Nano-Bioengineering, Graduate School of Incheon National University, Incheon 22012, Republic of Korea; Research Group of Gut Microbiome, Korea Food Research Institute, Wanju 55365, Republic of Korea; Division of Bioengineering, Incheon National University, Incheon 22012, Republic of Korea

**Keywords:** *Bacillus velezensis*, biofilm, 1-deoxynojirimycin, glucosyltransferase, *Streptococcus mutans*

## Abstract

*Streptococcus mutans* plays a key role in the development of dental caries and promotes the formation of oral biofilm produced by glucosyltransferases (GTFs). *Bacillus velezensis* K68 was isolated from traditional fermented foods and inhibits biofilm formation mediated by *S. mutans*. Gene amplification results demonstrated that *B. velezensis* K68 contained genes for the biosynthesis of 1-deoxynojirimycin (1-DNJ), a known GTF expression inhibitor. The presence of the GabT1, Yktc1, and GutB1 genes required for 1-DNJ synthesis in *B. velezensis* K68 was confirmed. Supernatant from *B. velezensis* K68 culture medium inhibited biofilm formation by 84% when *S. mutans* was cultured for 48 h, and inhibited it maximally when 1% glucose was added to the *S. mutans* culture medium as a GTF substrate. In addition, supernatant from *B. velezensis* K68 medium containing 3 ppb 1- DNJ decreased *S. mutans* cell surface hydrophobicity by 79.0 ± 0.8% compared with that of untreated control. The supernatant containing 1-DNJ decreased *S. mutans* adherence by 99.97% and 98.83% under sugar-dependent and sugar-independent conditions, respectively. *S. mutans* treated with the supernatant exhibited significantly reduced expression of the essential GTF genes *gtfB*, *gtfC,* and *gtfD* compared to that in the untreated group. Thus, *B. velezensis* inhibits the biofilm formation, adhesion, and GTF gene expression of *S. mutans* through 1- DNJ production.

**IMPORTANCE:** Dental caries is among the most common infectious diseases worldwide, and its development is closely associated with physiological factors of bacteria, such as the biofilm formation and glucosyltransferase production of *Streptococcus mutans.* Biofilms are difficult to remove once they have formed due to the exopolysaccharide matrix produced by the microorganisms residing in them; thus, inhibiting biofilm formation is a current focal point of research into prevention of dental caries. This study describes the inhibitory properties of *Bacillus velezensis* K68, an organism isolated from traditional Korean fermented foods, against biofilm formation by *S. mutans*. Herein, we show that *B. velezensis* inhibits the biofilm formation, adherence to surfaces, and glucosyltransferase production of *S. mutans*.

## INTRODUCTION

Dental caries is among the world’s major diseases, presenting in 60–90% of schoolchildren and adults, and is caused by microorganisms using various dietary saccharides, such as glucose, fructose, and sucrose (1). Especially, physiological properties such as acid tolerance, biofilm formation, and expression of virulence genes, of Streptococci in the mouth produced by the quorum-sensing system are known to affect oral health (2). *Streptococcus mutans* is a gram-positive, facultative anaerobic bacterium and is the primary causative agent of cariogenicity (3). *S. mutans* forms a biofilm and attaches to tooth surfaces, and the hydrophobicity of the biofilm is related to surface proteins antigens such as antigen I/II (SpaP), WapA, and SloC, on *S. mutans* (4). Oral biofilm produced by *S. mutans* is formed by microbial communities attached to the enamel layer of tooth surfaces (5, 6).

Biofilm is not easily removed because it is enclosed in a matrix of polysaccharides; thus, inhibition of the initial stages of biofilm formation is an important area of study (7). Biofilm is synthesized by glucosyltransferases (GTFs), which produce extracellular polysaccharides, in *S. mutans* (8). GtfB synthesizes α-1,3-linked insoluble glucan; GtfC synthesizes both insoluble and soluble glucan; and GtfD synthesizes α-1,6-linked soluble glucan (9–11). Various studies have been conducted on the anti-caries effects of microorganisms that inhibit the growth of *S. mutans* using treated culture medium from lactic acid bacteria and antibiotics or lytic enzymes from *Bacillus* spp. (12–17). Quercitrin and deoxynojirimycin (DNJ) were recently shown to inhibit the GTF gene expression in *S. mutans*, which led to biofilm formation (18).

This study focused on inhibition of *S. mutans* oral biofilm by a *Bacillus velezensis* strain. *B. velezensis* K68 was isolated from traditional fermented foods, and its inhibitory activity against *S. mutans* biofilm was analyzed. Additionally, the effects of treatment with *B. velezensis* culture supernatant were determined on *S. mutans* cell surface hydrophobicity, adherence, and regulation of virulence genes. We demonstrated that *B. velezensis* may prevent dental caries through the inhibition of biofilm produced by *S. mutans.*

## RESULTS AND DISCUSSION

### *S. mutans* biofilm inhibition by *B. velezensis* K68 isolated from fermented food

Because metabolites from some *Bacillus* species are known to inhibit biofilm formation by *S. mutans*, we isolated a *Bacillus* strain K68 exhibiting inhibitory effects on biofilms from fermented food. The 16S rRNA gene sequence (GenBank accession no. MG589484) of the isolated strain K68 was 99.64% similar to that of *Bacillus velezensis* CR-502 (Fig. S1). To investigate the inhibition of biofilm formation by *S. mutans* KCTC 3065, cells were treated with the supernatant from *B. velezensis* K68 culture medium containing 1% glucose and incubated at 37°C for different time periods (Fig. 1). In general, biofilm formation time for *S. mutans* in medium with 1% glucose is 24–48 h. The weak inhibition at the initial time (6 h) is likely due to insufficient biofilm formation at the time. After 12 h, *B. velezensis* K68 culture significantly inhibited biofilm formation compared to negative control (*B. subtilis* 142). In addition, the inhibitory effect increased in a concentration-dependent manner with *B. velezensis* K68 culture medium supernatant. Lytic enzymes and antibiotics produced by *Bacillus* spp. inhibit the growth of *S. mutans*, preventing biofilm formation (16, 17). Interestingly, there was no significant effect on *S. mutans* growth after treatment with *B. velezensis* K68 culture medium (data not shown). Previous studies showed that 1- deoxynojirimycin (1-DNJ) from mulberry (*Morus alba*) prevents biofilm formation and adhesion of *S. mutans* (7). Some *Streptomyces* and *Bacillus* spp. are known to produce 1-DNJ (19, 20). Therefore, *B. velezensis* K68 presumably inhibits the biofilm formation of *S. mutans* by producing 1-DNJ.

**FIG 1.**
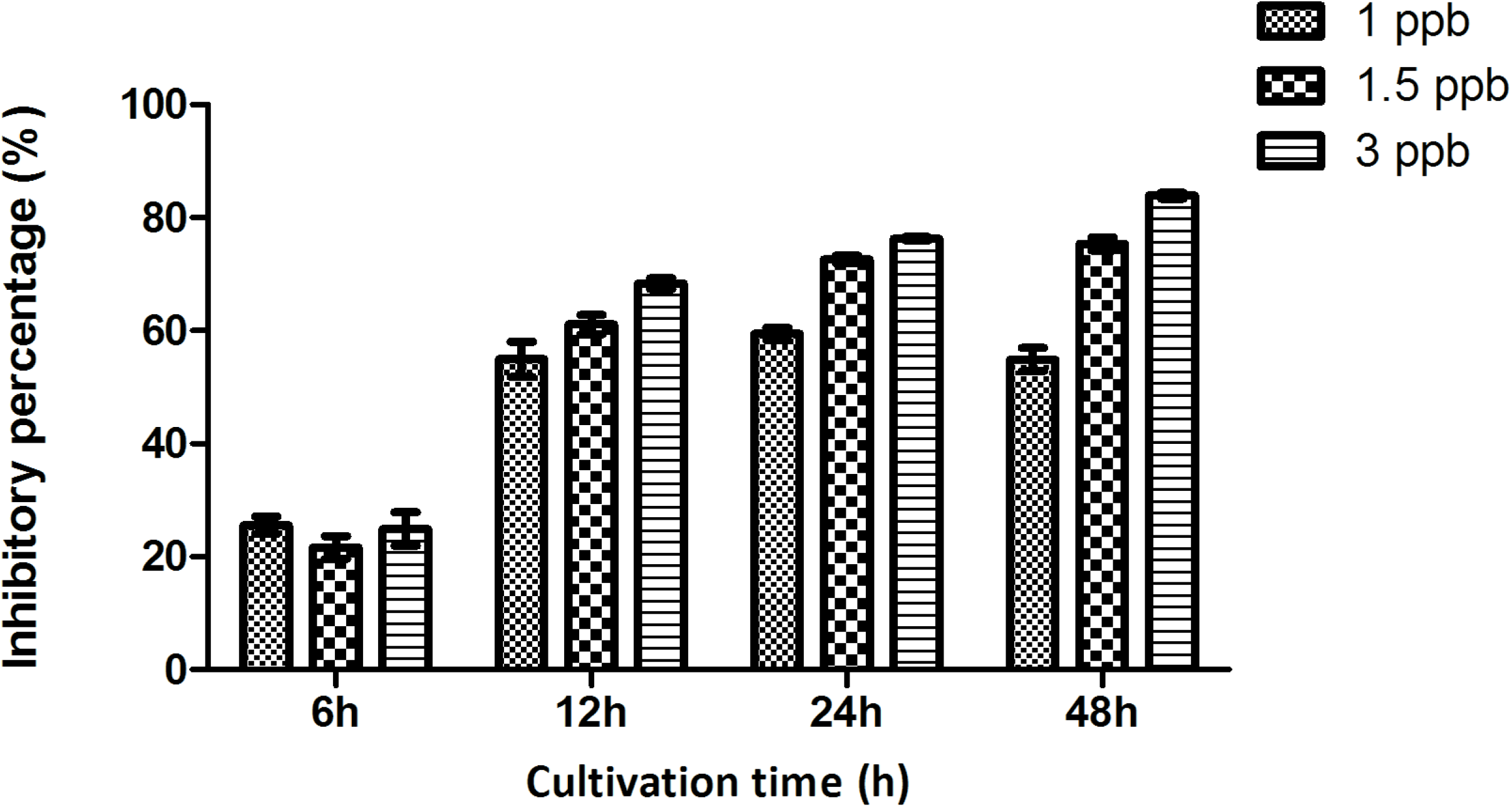
Inhibition of biofilm formation with supernatant from *B. velezensis* K68 culture media.

### Genetic and analytical confirmation of 1-DNJ production by *B. velezensis* K68

Some *Bacillus* spp. produce 1-DNJ and a promising operon including GabT1 (putative aminotransferase), Yktc1 (putative phosphatase), and GutB1 (putative oxidoreductase) could be essential to its biosynthesis (21). To determine whether *B. velezensis* K68 produces 1-DNJ, the presence of these three genes was confirmed by PCR (Fig. S2). A 1.2 kb fragment of *gabT1*, 0.9 kb of *yktc1*, and 1.0 kb of *gutB1* were successfully amplified. The obtained genes showed sequence differences from that of known 1-DNJ biosynthetic genes. The deduced amino acid sequence of each gene was analyzed, resulting that the GabT1, Yktc1, and GutB1 of *B. velezensis* K68 showed the highest homology to each protein from *B. velezensis* FZB42 (Table 1). The putative 1-DNJ biosynthetic gene cluster sequence assembled with *gabT1*, *yktc1* and *gutB1* of *B. velezensis* K68 has been deposited in GenBank under accession no. MH142722.

Triple quadrupole LC-mass spectrometry analysis showed that *B. velezensis* K68 produced 1-DNJ (data not shown). ESI in positive ion mode resulted in a peak on the chromatogram at 164.16 *m/z* for [M+H]^+^. This peak was confirmed to represent 1-DNJ (C_6_H_13_NO_4_, 163.17 g/mol) with a retention time of 9.19 min. The 1-DNJ in the supernatant of the *B. velezensis* K68 culture medium was identical to a standard 1-DNJ solution, suggesting that *B. velezensis* K68 produced 1-DNJ.

**Table 1.** Comparison of the deduced amino acid sequences of the 1-DNJ biosynthetic genes isolated from *B. velezensis* K68

| *B. velezensis* K68<br>(Nucleotide accession no.,<br>nucleotides, aa<sup>a</sup>) | *B. velezensis* FZB42<sup>b</sup><br>(aa, % identity/similarity to<br>K68) | Predicted function |
| --- | --- | --- |
| *gabT1* (MH142719, 1269, 422) | ABS72608 (425, 99.1%/99.1%) | 4-aminobutyrate<br>aminotransferase |
| *yktc1* (MH142720, 951, 316) | ABS72609 (316, 99.1%/99.4%) | fructose-1,6<br>biphosphate/inositol<br>monophosphatase |
| *gutB1* (MH142721, 1047, 348) | ABS72610 (348, 99.4%/99.7%) | zinc-binding<br>dehydrogenase |

<sup>a</sup>Deduced amino acid sequences for each gene of *B. velezensis* K68.
<sup>b</sup>Amino acid accession no. for each ORF of *B. velezensis* FZB42.

### Inhibitory effect of *B. velezensis* K68 on hydrophobicity and adherence of *S. mutans*

Cell surface hydrophobicity is involved in interactions between bacterial and epithelial cells and is important to initial bacterial adherence to tooth surfaces (22, 23). After treatment with 3 ppb 1-DNJ from the supernatant of *B. velezensis* K68 culture medium, *S*. *mutans* cell-surface hydrophobicity dramatically decreased from 79.0 ± 0.8% (untreated) to 8.1 ± 2.1% (treated) (Fig. 2A). The cell-surface hydrophobicity of *S. mutans* is known to be related to its cell-surface proteins. Hasan *et al* argued that quercitrin and 1-DNJ reduced *S. mutans* hydrophobicity by binding proteins on its cell surface (Ag I / II) (18). Its hydrophobicity was likely reduced by the 1-DNJ present in the supernatant of the *B. velezensis* K68 culture medium.

**FIG 2.**
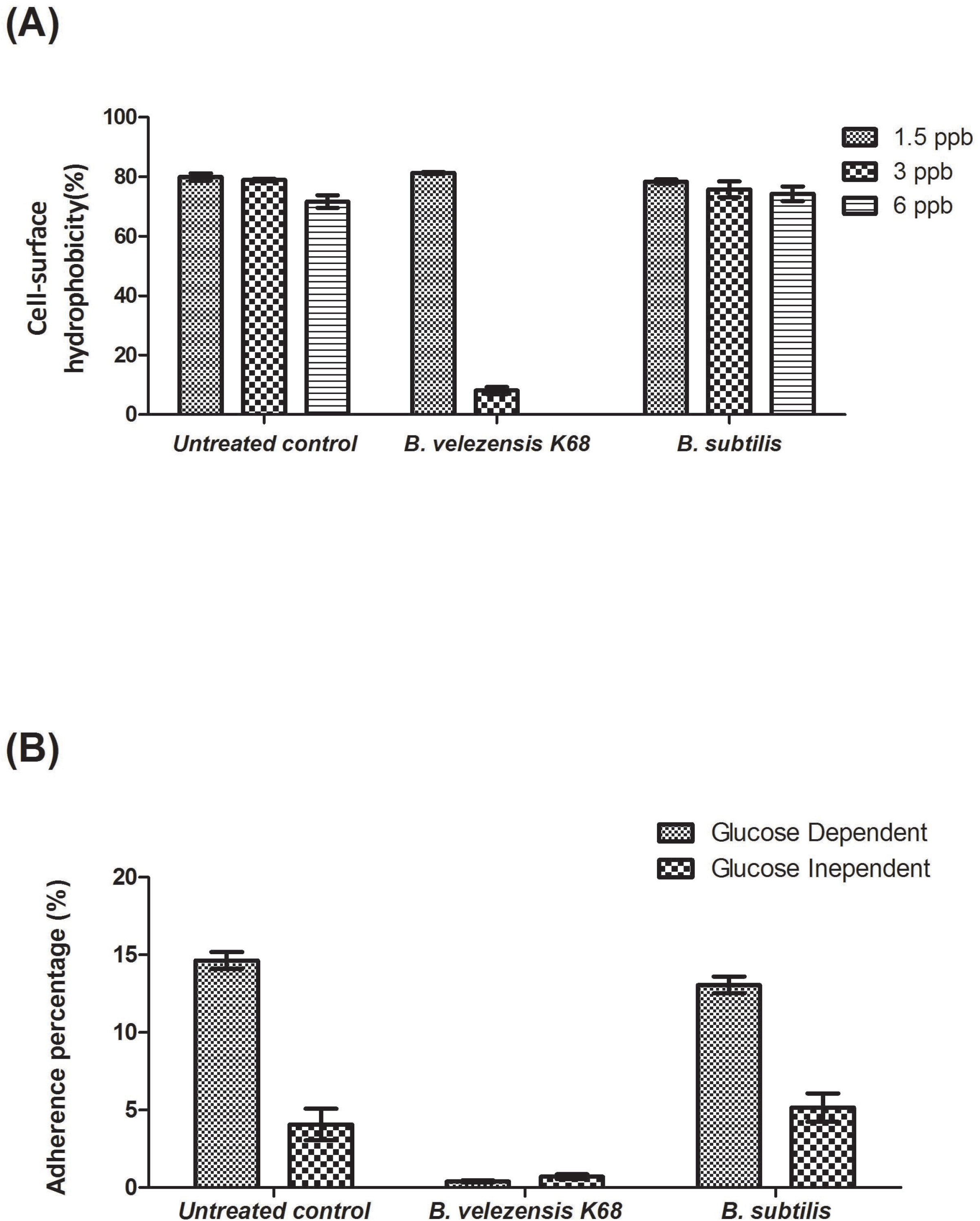
Effects of the supernatant from *B. velezensis* K68 culture medium on the cell-surface hydrophobicity (A) and adherence (B) of *S. mutans*.

Adhesion of *S. mutans* occurs by sugar-dependent and sugar-independent mechanisms. Sugar-dependent attachment is mediated by glucan produced from sugars through glucosyltransferases, whereas sugar-independent attachment is mediated by physicochemical forces, such as electrostatic forces, hydrophobic interactions, and hydrogen bonding with the constituents of saliva (24). In this study, the addition of glucose improved the adhesion of *S. mutans* (Fig. 2B). However, *B. velezensis* K68 culture supernatant containing 3 ppb 1-DNJ induced 99.97% and 98.83% decreases in adherence under sugar-dependent and sugar-independent conditions, respectively, as compared to the adherence of untreated control. Previous studies showed that 1-DNJ reduced the adherence of *S. mutans* in both sugar-dependent and sugar-independent conditions and showed a greater effect in the presence of sugar (18). *S. mutans* produces water-soluble glucans, such as dextran, and water-insoluble glucans, such as mutan, from sugars using various glucosyltransferases (25, 26). 1-DNJ is known to be an expression inhibitor of glucosyltransferases (18). Thus, our findings indicate that *B. velezensis* K68 produces 1-DNJ, which binds to cell surface proteins on *S. mutans* and acts as a competitive inhibitor of glucosyltransferases to reduce the hydrophobicity and adhesion of *S. mutans*.

### Inhibition of exopolysaccharide synthesis by *B. velezensis* K68

Scanning electron microscopy (SEM) revealed the formation of exopolysaccharides synthesized by *S. mutans* (18). The scanning electron micrographs show the effects of *B. velezensis* K68 culture medium supernatant on the ability of *S. mutans* to synthesize extracellular polysaccharides. The sample treated with the supernatant (Fig. 3B) showed significant dispersion of its cells, suggesting a reduction in exopolysaccharide synthesis. In contrast, the untreated and negative control samples (Fig. 3A, 3C) showed clear aggregation of cells immobilized in the exopolysaccharide pool. These results are consistent with 1-DNJ treatment of *S. mutans* (18). The effects of 1-DNJ are thought to be due to inhibition of the glucan synthesis, attachment, and biofilm formation of *S. mutans*.

**FIG 3.**
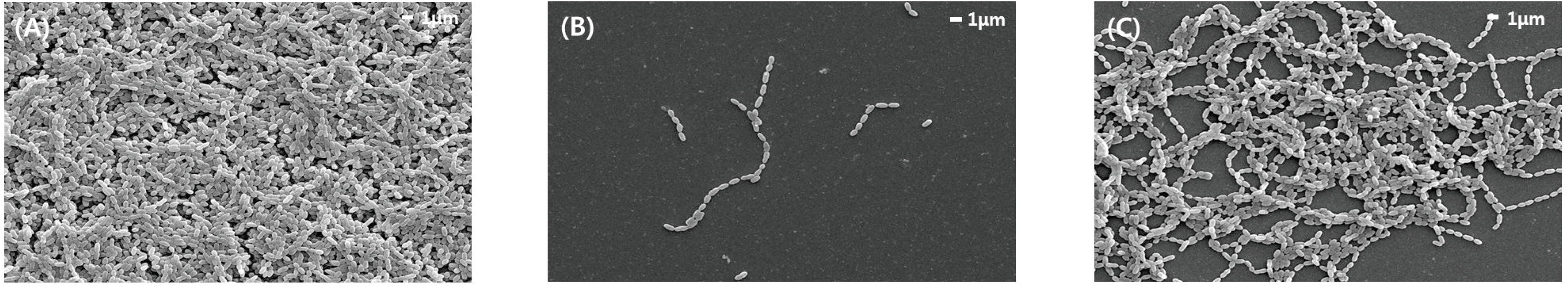
Scanning electron micrographs of *S. mutans* biofilms. (a) *S. mutans* (untreated control), (b) *S. mutans* treated with supernatant from *B. velezensis* K68 medium, (c) *S. mutans* treated with supernatant from *B. subtilis* medium (negative control).

### Inhibitory effect of *B. velezensis* K68 on glucosyltransferase expression in *S. mutans*

Three glucosyltransferase genes, *gtfB*, *gtfC*, and *gtfD*, are involved in water-soluble or water-insoluble glucan synthesis in *S. mutans* (27). The mRNA expressions of these genes were significantly decreased after treatment with *B. velezensis* K68 culture medium supernatant compared to their levels in untreated and negative control groups (Fig. 4). The *gtfB* and *gtfC* produce water-insoluble glucans, which act as adhesion molecules to immobilize bacteria on tooth pellicles (9). Therefore, *gtfB* and *gtfC* are critical toxic factors related to cariogenicity (8). Both genes were independently expressed, and there was a difference between their promoters (28). Additionally, the expression of the two genes is decreased by ions (Ca, K, and Mg) (29). Interestingly, *gtfB* mRNA levels decreased in the *B. subtilis*-treated group, likely due to the presents of ions in the *B. subtilis* culture medium. In the *B. velezensis* K68-treated group, the mRNA expression of all three *gtf* genes was significantly reduced, and thus the biofilm formation and adherence of *S. mutans* were decreased compared to those in the untreated group. These results were consistent with those observed in treatments with *Lactobacillus salivarius* and 1-DNJ (14).

**FIG 4.**
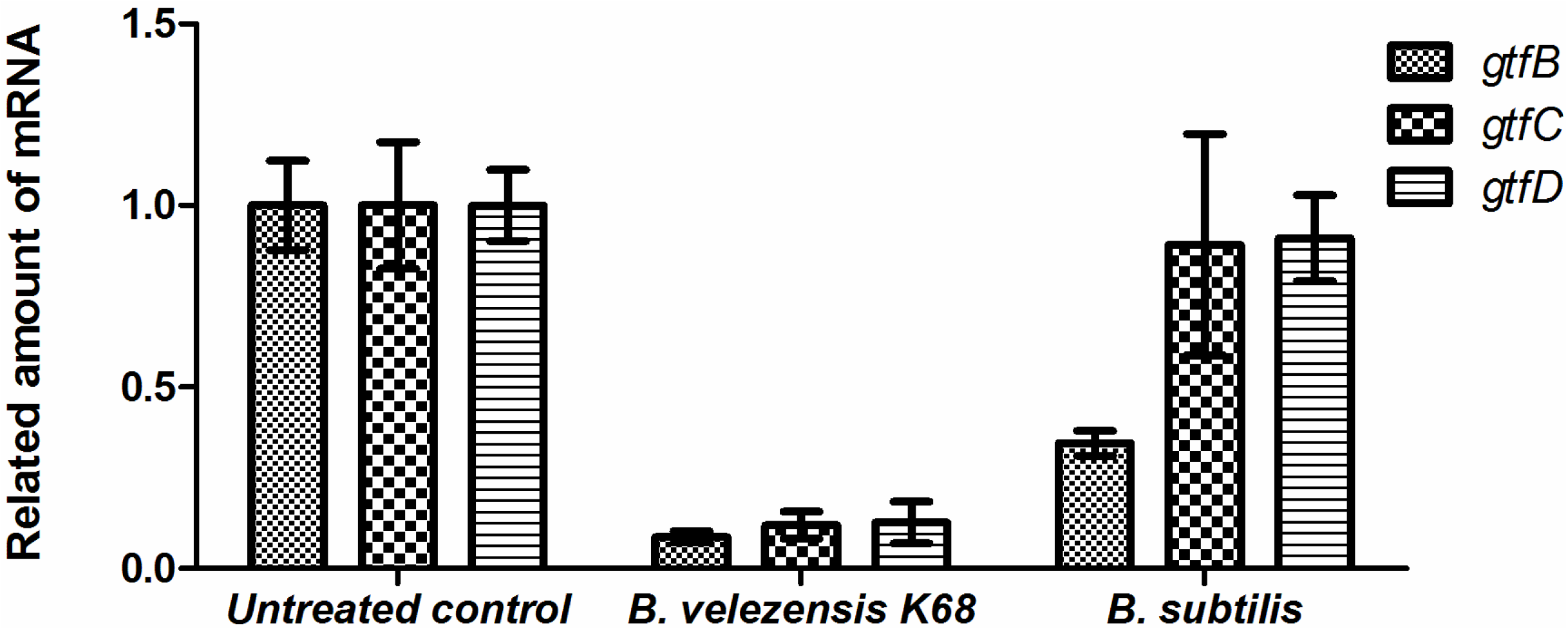
*S. mutans* glucosyltransferase gene expression after treatment with supernatant from *B. velezensis* K68 medium. *gtfB*, *gtfC*, and *gtfD*, which encode glucosyltransferases in *S. mutans*, were analyzed by qPCR.

This study was conducted to determine the ability of *B. velezensis* K68, which was isolated from traditional Korean fermented foods, to inhibit biofilm formation by *S. mutans*. The 1-DNJ biosynthetic gene in *B. velezensis* K68 was successfully amplified through PCR, and 1-DNJ production was confirmed by LC-Mass. The 1-DNJ produced by *B. velezensis* K68 inhibited the expression of the GTF genes responsible for biofilm formation by *S. mutans*. In addition, physicochemical analyses showed that culture medium containing 1-DNJ inhibited biofilm formation by *S. mutans*. These results indicate that *B. velezensis* K68 can be used to produce fermented food useful for dental health. The applicability of *B. velezensis* K68 warrants further investigation.

## MATERIALS AND METHODS

### Bacterial strains and culture conditions

*Streptococcus mutans* KCTC 3065 was purchased from the Korean Collection for Type Cultures (Daejeon, Republic of Korea). *Bacillus velezensis* K68 was isolated from traditional Korean fermented foods. *S. mutans* KCTC 3065 was routinely cultured in brain-heart infusion (BHI) broth (MB Cell, Los Angeles, CA, USA), and *B. velezensis* K68 was successively cultured in tryptic soy broth (TSB; BD Biosciences, Franklin Lakes, NJ, USA) at 37°C with continuous shaking for 3 d. After cultivation, these media were centrifuged at 10,000 rpm for 10 min, and the supernatant was filtered with a 0.2 μm filter to remove cell debris.

### Isolation of *Bacillus* strains harboring 1-DNJ biosynthetic genes

Extraction of genomic DNA from *B. velezensis* K68 was performed using phenol-chloroform isoamylalcohol as previously described (30). The 16S rRNA and 1-DNJ biosynthesis genes of *B. velezensis* K68 were amplified by polymerase chain reaction (PCR). The PCR primer sets for the genes were as follows: 27F (5′-AGA GTT TGA TCM TGG CTC AG-3′) and 1492R (5′-CGG TTA CCT TGT TAC GAC TT-3′) for the 16S RNA gene, MJS-23 (5′-ATG GGA ACG AAG GAA ATC ACG AAT CCA-3′) and MJS-24 (5′-TCA CTT GAT TTC CTC CAA TAG CTT GCG-3′) for *gabT1*, MJS-19 (5′-GTG AGA GAC TAT ATC ATY GRG CTT GGA-3′) and MJS-20 (5′-TTA GGA GTC CAG ACC AAC GCC TTC ATA-3′) for *yktc1*, and MJS-21 (5′- ATG AAG GCG TTG GTC TGG ACT CCT AAT-3′) and MJS-22 (5′-TTA TAA AAG TTY CGG ATC AGA CAC RAG-3′) for *gutB1* (31). PCR was performed at 95°C for 5 min, followed by 30 cycles at 95°C for 30 s, 55°C for 30 s, 72°C for 1 min 30 s, and then a final extension at 72°C for 7 min. The PCR products were sequenced and analyzed for the construction of a phylogenetic tree. The samples were dissolved in LC/MS-grade methanol (Thermo Fisher Scientific, Waltham, MA, USA). Triple quadrupole LC analysis was conducted with a Finnigan TSQ Quantum Ultra EMR, a TSK gel amide-80 column (3 μm, 2.0 × 150 mm; Tosoh, Tokyo, Japan), and 10 μL of injected sample volume. Chromatograms were obtained using a flow late of 0.2 mL/min with 5 mM ammonium acetate as eluent A and acetonitrile as eluent B. Elution gradient conditions were set at 0 min (90% B), 10 min (40% B), 12 min (40% B), 13 min (90% B), and 20 min (90% B).

### Biofilm formation assay

Biofilm formation was assessed using flat-bottomed, polystyrene, 96-well microtiter plates. *S. mutans* KCTC 3065 was cultivated overnight in BHI broth, and the resulting cell suspension was diluted to 0.8−1.0 optical density (OD) at 600 nm. A total of 250 μL of liquid, composed of 50 μL of the cell suspension, 150 μL of BHI broth containing 1% glucose, and 50 μL of *B. velezensis* K68 culture supernatant, was injected into each well of the microtiter plates. The liquid was mixed and incubated at 37°C for 6, 12, 24, and 48 h, and planktonic cells were gently removed with sterile water. Biofilm formation was compared with that of *Bacillus subtilis* as a negative control. Attached cells in the wells were fixed with formalin (37%, diluted 1:10) containing 2% sodium acetate for 1 h. Each well was stained with 250 μL of 0.1% Crystal violet for 15 min at 25°C and rinsed with sterile water. Bound dye was removed with 150 μL of 95% ethanol. Biofilm formation was observed by measuring the suspension at 595 nm with a microplate reader (Multiskan FC plate reader; Thermo Fisher Scientific) (18). Inhibition percentage was calculated as follows: inhibition percentage = (untreated sample OD_595_ – test sample OD_595_)/untreated sample OD_595_ × 100.

### Triple quadrupole LC-mass spectrometry

Triple quadrupole LC-mass spectrometry was used to confirm 1-DNJ production by *B. velezensis* K68. Cultivated samples were centrifuged at 10,000 rpm for 10 min (1580R, Labogene, Seoul, Korea), and the supernatant was filtered with a 0.2 μm filter and freeze-dried (TFD8503; Ilshinbiobase, Seoul, Korea) for liquid chromatography-mass spectrometry confirmation of 1-DNJ synthesis. The samples were dissolved in methanol (LC/MS Grade, Thermo Fisher Scientific). Triple quadrupole LC analysis was conducted with a Finnigan TSQ Quantum Ultra EMR system and TSK gel amide-80 column (3 μm, 2.0× 150 mm; Tosoh) injected with 10 μL samples. The chromatogram was obtained using a flow late of 0.2 mL/min with 5mM ammonium acetate as eluent A and acetonitrile as eluent B. Different elution gradient conditions were set for 0 min (90% B), 10 min (40% B), 12 min (40% B), 13 min (90% B), and 20 min (90% B).

### Scanning electron microscopy

Mature *S. mutans* KCTC 3065 was grown at 37°C for 24 h in BHI containing 1% (w/v) glucose and placed on glass coverslips in cell culture plates (60 mm × 15 mm). The coverslips were gently washed twice with 1× phosphate-buffered saline (PBS). Adherent cells were fixed with PBS containing 2.5% glutaraldehyde for 1 h and gradually dehydrated with a series of 25, 50, 75, 90, and 100% ethanol treatments. The prepared cells were dried completely in an oven for 1 h. The cells were then coated with gold, and their morphologies observed using an JSM-7001F field emission scanning electron microscope (JEOL Korea, Seoul, Republic of Korea) (32).

### Cell-surface hydrophobicity

The hydrophobicity of the *S. mutans* KCTC 3065 cell surface was measured by comparing percentages of binding affinity of the cells to toluene. Cells cultivated in BHI broth were washed twice with 0.85% (w/v) NaCl solution. They were then resuspended to an approximate OD_600_ of 0.3. To calculate hydrophobicity, 3 mL of cell suspension was added to 250 μL of toluene. This solution was mixed with a vortex for 2 min and incubated at 25°C to allow phase separation. After separation of the toluene phase, the OD of the aqueous phase at 600 nm was recorded with a spectrophotometer (33). The hydrophobicity percentage was calculated as follows: (OD _initial_ – OD _final_) / OD _initial_ × 100.

### Bacterial adherence assay

The ability of *S. mutans* KCTC 3065 to adhere to glass surfaces was determined as previously described (34). Cells were grown at 37°C for 24 h in glass test tubes at an angle of 30° with BHI broth containing 1% (w/v) glucose. After incubation, the tubes were gently spun down, and then the planktonic cells were removed. The attached cells were suspended in 0.5 M NaOH by vortexing and quantified at OD_600_. The adherent percentage was calculated as follows: (OD_adherent cells_/OD_total cells_) × 100.

### Quantitative real-time polymerase chain reaction

Total RNA was extracted from *S. mutans* KCTC 3065 cultures treated with *B. velezensis* K68 culture supernatant using RNeasy mini-columns (Qiagen, Hilden, Germany) with proteinase K solution (Qiagen) (35). cDNA was synthesized with a DiaStar RT kit (Solgent, Daejeon, Korea) using random primers. Quantitative real-time reverse transcription polymerase chain reaction (qRT-PCR) was performed with 2× Real-Time PCR mix (including SYBR green; Solgent) using a StepOnePlus Real-Time PCR System (Applied Biosystems, Foster City, CA, USA) run at 95°C for 5 min, followed by 40 cycles at 95°C for 15 s, 60°C for 1 min, and then an annealing and extension step at 60°C for 5 min. Primer sequences were as follows: 5′-AGC AAT GCA GCC ART CTA CAA AT-3′ and 5′-ACG AAC TTT GCC GTT ATT GTC A-3′ for *gtfB*, 5′-YCT CAA CCA ACC GCC ACT GTT-3′ and 5′- TTA ACG TCA AAA TTA CGA CAT AAT C-3′ for *gtfC*, 5′-CAC AGG CAA AAG CTG AAT TAA CA-3′ and 5′-AAT GGC CGC TAA GTC AAC AG-3′ for *gtfD*, and 5′-CCT ACG GGA GGC AGC AGT AG-3′ and 5′-CAA CAG AGC TTT ACG ATC CGA AA-3′ for the 16S rRNA gene. Relative quantification was performed using the 2-ΔΔCt method with a reference gene (16S rRNA gene) from *S. mutans* KCTC 3065 (36, 37).

## SUPPLEMENTAL MATERIAL

Supplemental material for this article may be found.

**SUPPLEMENTAL FILE 1,** PDF file, 0.2 MB.

## ACKNOWLEDGMENTS

This research was supported by the Basic Science Research Programs through the National Research Foundation of Korea (NRF) funded by the Ministry of Science, ICT & Future Planning (NRF-2014R1A1A1002980) and the Ministry of Education (NRF-2016R1D1A1B03931582) for M.-J.S. This work was also supported by the Main Research Program (E0170602-02) of the Korea Food Research Institute (KFRI) funded by the Ministry of Science and ICT & Future Planning for Y.-D.N.

